# Decoding *Klebsiella pneumoniae* in Poultry Chain: Unveiling Genetic Landscape, Antibiotic Resistance, and Biocide Tolerance in Non-Clinical Reservoirs

**DOI:** 10.1101/2024.01.23.576544

**Authors:** Joana Mourão, Mafalda Magalhães, Marisa Ribeiro-Almeida, Andreia Rebelo, Carla Novais, Luísa Peixe, Ângela Novais, Patrícia Antunes

## Abstract

The rise of antibiotic resistance in the food chain is influenced by the use of antimicrobial agents, such as antibiotics, metals, and biocides, throughout the entire farm-to-fork continuum. Besides, non-clinical reservoirs potentially contribute to the transmission of critical pathogens such as multidrug-resistant (MDR) *Klebsiella pneumoniae*. However, limited knowledge exists about the population structure and genomic diversity of *K. pneumoniae* circulating in conventional poultry production. We conducted a comprehensive characterization of *K. pneumoniae* across the whole chicken production chain (flocks/environment/meat, 2019-2022), exploring factors beyond antibiotics, like copper and quaternary ammonium compounds (QACs). Clonal diversity and adaptive features of *K. pneumoniae* were characterized through cultural, molecular (FT-IR), and whole-genome-sequencing (WGS) approaches. All except one flock were positive for *K. pneumoniae* with a significant increase (p < 0.05) from early to pre-slaughter stages, most persisting in chicken meat batches. Colistin-resistant *K. pneumoniae* rates were low (4%), while most samples carried MDR strains (67%) and copper-tolerant isolates (63%; *sil*+*pco* clusters; MIC_CuSO4_≥16mM), particularly at pre-slaughter. Benzalkonium chloride consistently exhibited activity in *K. pneumoniae* (MIC/MBC range=4-64mg/L) from diverse and representative strains independently of the presence/absence of genes linked to QACs tolerance. A polyclonal *K. pneumoniae* population, discriminated by FT-IR and WGS, included various lineages dispersed throughout the chicken’s lifecycle at the farm (ST29-KL124, ST11-KL106, ST15-KL19, ST1228-KL38), until the meat (ST1-KL19, ST11-KL111, ST6405-KL109, and ST6406-CG147-KL111), or over years (ST631-49 KL109, ST6651-KL107, ST6406-CG147-KL111). Notably, some lineages were identical to those from human clinical isolates. WGS also revealed F-type multireplicon plasmids carrying *sil*+*pco* (copper) co-located with *qacE*Δ1±*qacF* (QACs) and antibiotic resistance genes like those disseminated in humans. In conclusion, chicken farms and their derived meat are significant reservoirs for diverse *K. pneumoniae* clones enriched in antibiotic resistance and metal tolerance genes, some exhibiting genetic similarities with human clinical strains. Further research is imperative to unravel the factors influencing *K. pneumoniae* persistence and dissemination within poultry production, contributing to improved food safety risk management. This study underscores the significance of understanding the interplay between antimicrobial control strategies and non-clinical sources to effectively address the spread of antimicrobial resistance.

## 1 Introduction

Intensive poultry production is a crucial sector of the global food industry. It faces significant challenges in addressing the growing demand for poultry products, especially antibiotic-free chicken (Mottet and Tempio, 2017; Karcher and Mench, 2018). Ensuring biosecurity, which includes proper hygiene practices, vaccination programs, regular monitoring of flock health and effective farm management strategies, is essential to prevent disease transmission between flocks and farms. However, despite these efforts, intensive chicken production relies heavily on antibiotics and coccidiostats (Karcher and Mench, 2018; Gržinić et al., 2023). This reliance becomes even more concerning when considering that poultry presents potential risks to human health as it can be a reservoir of zoonotic pathogens causing infectious diseases and can contribute to the spread of antimicrobial resistance (AMR) within the food chain (Golden et al., 2021; European Food Safety Authority (EFSA) and European Centre for Disease Prevention and Control (ECDC), 2023). While antibiotics have traditionally been seen as the main drivers of AMR, recent restrictions on their use in food-producing animals, as well as alternative antimicrobial approaches, suggest the contribution of other compounds as selectors of antibiotic-resistant bacteria (Rebelo et al., 2023).

Metals and biocides are used in food-animal production for various purposes, including as feed additives, growth promoters, antiseptics, and disinfectants, to decrease the dependence on antibiotics [(EMA Committee for Medicinal Products for Veterinary Use (CVMP) and EFSA Panel on Biological Hazards (BIOHAZ) et al., 2017; Rebelo et al., 2023)]. Copper is one of the metals commonly added to chicken feed. Its antimicrobial properties not only improve animal nutrition and productivity (e.g., by modulating gut microbiota) but also reduce disease risks, thereby boosting the overall flock health (Broom et al., 2021; El Sabry et al., 2021). Furthermore, biocides formulated with quaternary ammonium compounds (QACs) are used to disinfect surfaces, equipment, feeding systems, and water sources. Their versatility and broad-spectrum antimicrobial activity assist in controlling and preventing pathogens dissemination between flocks within poultry facilities (Chen et al., 2023). Genes associated with tolerance to metals and QACs often share the same genetic contexts with antibiotic-resistance genes (Slifierz et al., 2015; Li et al., 2022; Pereira et al., 2023). Thus, the use of copper-supplemented feed and QAC-based biocides could contribute to co-selection effects (Webber et al., 2015; Kampf, 2018; Rebelo et al., 2023).

Current intensive chicken production involves large-scale operations that span breeding and hatching to rearing, processing, and distribution. However, the impact of diverse antimicrobial strategies used throughout farm to fork on the spread of AMR remains underexplored [(EMA Committee for Medicinal Products for Veterinary Use (CVMP) and EFSA Panel on Biological Hazards (BIOHAZ) et al., 2017; Rebelo et al., 2023)]. To effectively address AMR, a One Health approach – which emphasizes coordinated efforts across the domains of animals, humans, and the environment – is essential. This approach is not only crucial for elucidating the origins of less common foodborne pathogens such as *K. pneumoniae* but also for understanding emergent reservoirs and vectors of AMR genes outside the clinical setting (Wyres and Holt, 2018). Our prior study unveiled a significant occurrence of copper tolerance and multidrug resistance among *K. pneumoniae* strains found in chicken flocks (Mourão et al., 2023). We also identified *K. pneumoniae* lineages and plasmids carrying *sil+pco* copper tolerance and variable antibiotic resistance genes, resembling those identified in human clinical isolates worldwide (Mourão et al., 2023). These findings highlight the urgent need to further investigate the factors, drivers, and sources that contribute to the selection and persistence of MDR *K. pneumoniae* within the poultry production chain. However, there has been limited research focusing on the early stages of chicken rearing (e.g., one-day-old chicks) and the in-house poultry environment, including cleaned poultry houses post-vacancy, where broilers are raised for 30-35 days before being sent to slaughter for meat production (Daehre et al., 2018; Zhai et al., 2020).

This study aims to provide a comprehensive analysis of the occurrence, diversity, and persistence of *K. pneumoniae* in the whole poultry production chain (from one-day-old chicks to chicken meat) between 2019 and 2022. Furthermore, we assessed the contribution of factors other than antibiotics (use of copper and quaternary compounds) as putative selective agents of AMR genes and bacteria that are clinically relevant. Identifying the transmission sources and pathways for MDR *K. pneumoniae* will pave the way for devising effective strategies to mitigate its dissemination, ensuring animal welfare, environmental sustainability, public health, and overall improved food safety.

## 2 Material and Methods

### 2.1 Sampling design at the chicken farm and slaughterhouse processing plant

Our sampling included seven Portuguese intensive-based chicken farms with conventional indoor and floor-raised production systems in compliance with EU legislation, as indicated by the operator (ADAS, 2016). Six similar farms (arbitrarily designated as A, B, C, E, G, and H) previously studied (Mourão et al., 2023) and one (I) recently restructured with modern grow-out poultry house facilities were selected. In all farms, colistin was banned since January 2018, while copper was routinely used as an additive in inorganic formulation feed (far below the maximum dose of 25 mg/kg of Cu, according to EU Regulation 2018/1039). The vacancy period varied from 11 to >15 days, being each poultry house depopulated, cleaned, and disinfected using similar routine cleaning and disinfection procedures. The biocide active compounds used were benzalkonium chloride (BZC) or didecyl dimethyl ammonium chloride (DDAC) and hydrogen peroxide for water distribution systems.

In each of the seven farms, female, and male mixed one-day-old chicks (Ross 308 strain) were randomly distributed by two poultry houses at arrival (around 5000-60.000 chickens per flock/house). A total of 14 flocks were followed and sampling was carried out over three different periods during the production cycle (the length ranged from 26-43 days) between February and May 2022 (**Figure 1**). The initial sampling period included collecting samples from one-day-old chicken transport boxes (P0; n=14 samples; each sample comprising 12 boxes). These were obtained using stick swabs, with each swab tube containing 10 ml of Buffered Peptone Water-BPW and used in 4 chicken boxes (a total of three swabs per sample). Additionally, the inside of cleaned poultry houses (P1; n=14 samples) were sampled using the boot swab technique, which employs dry-foot swabs in a zigzag pattern, according to EU Regulation 200/2012. In the second period, the inside of grow-out houses containing the same flocks the day before slaughter (P2; n=14 samples; after >25 days) was collected using the boot swab technique. In the third period, raw chicken meat samples (P3; n=14 batches) from the same flocks were collected after slaughter and air chilling at the poultry production slaughterhouses, immediately before distribution for retail sale. Each sample (50 g) was processed as a pool of neck skin from 10-15 carcasses of the same batch (each batch corresponded to one flock from the same farm and poultry house slaughtered at the same time) (according to EU Regulation 1086/2011).

**Figure 1.**
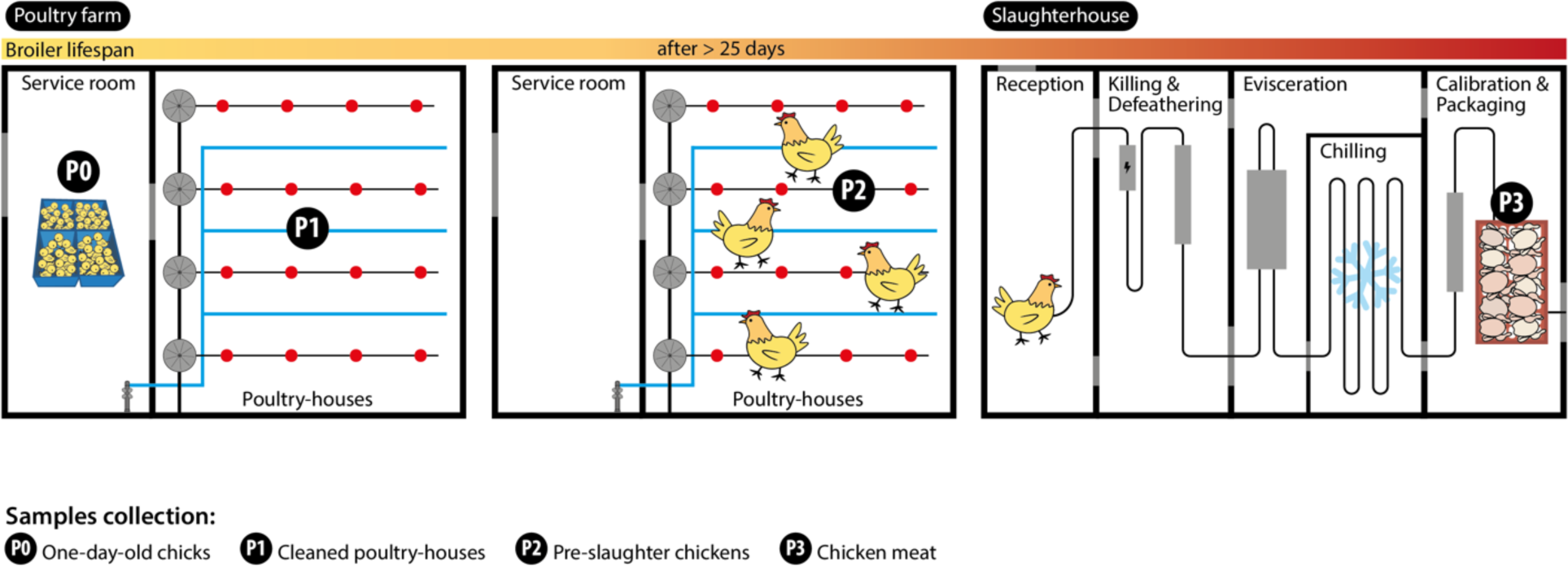
Sampling strategy at the chicken farm and slaughterhouse processing plant. Sample collection points are indicated by P0, P1, P2 and P3.

All previous samples were collected in sterile plastic bags/containers, transported at 4°C to the laboratory, and processed on the same day. Subsequent sample processing was performed using cultural/molecular approaches as described in the following sections.

### 2.2 Screening of Klebsiella pneumoniae

*K. pneumoniae* was recovered from Simmons citrate agar plates with 1% inositol (SCAi), the most suitable selective medium for *Klebsiella* recovery, directly from the suspended sample and after enrichment. A common initial step consisted of mixing swabs (three pooled swab tubes-P0 and two swab foot-P1 and P2) or weighing 25 g of pooled meat samples (P3) into 1/10 mL of BPW and BPW supplemented with 3.5 mg/L colistin. The direct culture method included spreading an aliquot of 100 µL of BPW and BPW+colistin after 1 h at room temperature (resuscitation step) on SCAi supplemented or not with colistin (3.5 mg/L). The enrichment approach involved the same procedure but after a previous incubation of BPW and BPW+colistin at 37°C for 16-18 h. All the SCAi plates were incubated at 37°C for 48h. One to five colonies of each presumptive morphotype were selected for identification. Isolates were identified presumptively by CHROMagar™ Orientation and then by PCR for *K. pneumoniae* (Bialek-Davenet et al., 2014).

### 2.3 *Klebsiella pneumoniae* diversity between samples

We used Fourier Transform Infrared (FT-IR) spectroscopy with attenuated total reflectance (ATR) to infer isolates’ relatedness between isolates identified in the same or different samples. After growth under standardized culture conditions (Mueller-Hinton agar; 37°C/18h), a colony was directly deposited and air-dried on the ATR accessory of the FT-IR instrument. Spectra were acquired in a Spectrum Two instrument (Perkin-Elmer, USA) under standard conditions (4000-600 cm^-1^, 4 cm-1 resolution, and 16 scan co-additions). The region between 1200cm^-1^ and 900 cm^-1^ was compared with each other, and with those from two different machine-learning classification models, used to predict *K. pneumoniae* capsular (KL) types: i) a Random Forest classification model that enables identification of up to 33 KL-types from well-characterized international *K. pneumoniae* clones from the clinical setting (Novais et al., 2023a); and ii) a Random Forest classification model to allow identification of up to 21 KL-types (eleven in common with the previous model) based on a spectral database of poultry isolates identified in previous studies (Mourão et al., 2023). Isolate relatedness and prediction of KL-types were inferred as described previously (Novais et al., 2023a). Isolates predicted to have the same KL-type were considered putatively related. FT-IR-based assignments were confirmed by PCR of the *wzi* gene and further sequencing at Eurofins Genomics (https://www.eurofinsgenomics.eu/) to infer K-type using BIGSdb (http://bigsdb.pasteur.fr/klebsiella/klebsiella.html) (Brisse et al., 2013).

### 2.4 Antimicrobial susceptibility to antibiotics

Susceptibility to 17 antibiotics (amoxicillin+clavulanic acid-30µg, amikacin-30µg, aztreonam-30µg, cefepime-30µg, cefotaxime-5µg, cefoxitin-30µg, ceftazidime-10µg, chloramphenicol-30µg, ciprofloxacin-5µg, gentamicin-10µg, kanamycin-30µg, meropenem-10µg, nalidixic acid-30µg, sulfamethoxazole-300µg, tetracycline-30µg, tobramycin-10µg, and trimethoprim-5µg) was determined using the disc diffusion method. The Minimum Inhibitory Concentration (MIC) of colistin was determined using the reference broth microdilution method (European Committee for Antimicrobial Susceptibility Testing, 2016). *Escherichia coli* ATCC 25922 was used as the control strain. The results of both assays were interpreted using the European Committee of Antimicrobial Susceptibility Testing (EUCAST) (European Committee on Antimicrobial Susceptibility Testing, 2022) and, when this was not possible, the Clinical and Laboratory Standards Institute (CLSI) guidelines (Clinical and Laboratory Standards Institute, 2022). Isolates categorised as “susceptible, increased exposure” (EUCAST guidelines) or “intermediate resistant” (CLSI guidelines) were classified as susceptible. Multidrug resistance (MDR) was considered when the isolates were resistant to three or more antibiotics from different families (in addition to ampicillin, to which all *K. pneumoniae* are intrinsically resistant). Screening of colistin resistance genes (*mcr-1-5* and *mcr-6-9*) was performed for all isolates using two multiplex PCR (Rebelo et al., 2018; Borowiak et al., 2020).

### 2.5 Antimicrobial susceptibility to copper

Copper-Cu susceptibility in representative isolates (different farms, flocks, and genomic backgrounds) was evaluated using the agar dilution method under anaerobic conditions (Mourão et al., 2016). Briefly, the MIC was determined using Mueller-Hinton 2 agar freshly prepared and supplemented with different copper sulphate (CuSO_4_) concentrations (0.5 to 36 mM) and a final adjustment to pH = 7.2. The plates were then inoculated with 0.001 mL suspension (10^7^ CFU/mL) of each isolate and incubated at 37°C under anaerobic conditions for 18h-20h. The MIC was identified as the lowest concentration where no visible growth was observed. Control strains included *Escherichia coli* ED8739 carrying the plasmid pRJ1004 with the *sil* and *pco* cluster (MIC_CuSO4_=16-20mM) and *Enterococcus lactis* BM4105RF without acquired copper tolerance genes (MIC_CuSO4_=2-4 mM) (Novais et al., 2023b). All *K. pneumoniae* isolates were screened for the *silA* copper tolerance gene using a PCR, given its strong association with the presence of an intact *sil* operon and a Cu tolerance phenotype (Mourão et al., 2016, 2023).

### 2.6 Antimicrobial susceptibility to benzalkonium chloride

The MIC and minimum bactericidal concentrations (MBC) of benzalkonium chloride (BZC) (CAS 68391-01-5, VWR) were determined for representative sequenced isolates (n=45; different clones, sample types, farms, years and the presence or absence of QAC tolerance genes) using the Mueller-Hinton broth microdilution method (pH=7.2; 37°C/20h) (Clinical and Laboratory Standards Institute, 2018). Briefly, a 96-well microtiter plate containing serial two-fold dilutions of BZC (concentration ranging from 0.125 to 128 mg/L) was used to assess the susceptibility of bacterial suspensions in log-phase growth (adjusted to reach a final inoculum of 5×10^5^ CFU/mL in each well) at 37°C for 20h. Microdilution panels were freshly prepared before each assay. The first concentration of BZC without visible growth was considered the MIC (Clinical and Laboratory Standards Institute, 2018).

To determine the MBC_BZC_, 10μl of each well without visible growth from the 96-well MIC plate were incubated onto brain heart infusion agar plates at 37°C for 24h, as defined by the CLSI (Clinical and Laboratory Standards Institute, 1999). The MBC was defined as the lowest QAC concentration where the colony count was equal to or less than the CLSI-specified rejection value, based on the final inoculum count of each well (Clinical and Laboratory Standards Institute, 1999). Each experiment was repeated three times and the MIC/MBC values corresponded to the mean of these determinations. To ensure assay reproducibility, the *Enterococcus faecalis* ATCC 29212 (without any known QACs tolerance genes) was included as a control strain (MIC_BZC_ and MBC_BZC_ varied between 1-4 mg/L) (Pereira et al., 2023).

### 2.7 Genomic analysis by whole-genome sequencing

We then aimed to elucidate the sources and persistence of specific clonal lineages and/or MDR plasmids along the chicken production. For that, we compared 68 genomes, including 48 sequenced *de novo* in the present study (n=31 from 2022 farms and n=17 from 2019-2020 farms) and 20 obtained in the previous study (Mourão et al., 2023), representing different farms, stages, timespans, and KL-types. Genomic DNA was extracted from the isolates using the Wizard Genomic DNA purification kit (Promega Corporation, Madison, WI) and the final concentration was measured with a Qubit 3.0 Fluorometer (Invitrogen, Thermo Fisher Scientific, USA). Subsequently, the isolates were sequenced using the Illumina NovaSeq 6000 S4 PE150 XP (Illumina, San Diego, CA, USA) at Eurofins Genomics (https://eurofinsgenomics.eu/). FastQC v0.11.9 (Andrews, 2010) and MultiQC v1.13.dev0 (Ewels et al., 2016) with default parameters were used for the quality control of the raw sequence data. If needed, the reads were filtered using BBDuk v39.01 (http://sourceforge.net/projects/bbmap/), followed by *de novo* assembly with SPAdes v3.15.5 (Bankevich et al., 2012) integrated within Unicycler v0.5.0 (Wick et al., 2017). QUAST v5.0.2 (Gurevich et al., 2013) was used for the quality evaluation of assemblies and CheckM v1.2.2 (Parks et al., 2015) and BUSCO v5.4.6 (Simão et al., 2015) for genome completeness assessment.

The assemblies were annotated using the RAST server (Aziz et al., 2008). Genomes’ assemblies were then submitted to Pathogenwatch v2.3.3 (https://pathogen.watch/) which provided information on the capsular polysaccharide (K) and lipopolysaccharide (O) locus types and serotypes using Kaptive (Wyres and Holt, 2016), but also multi-locus sequence type (MLST) (Diancourt et al., 2005) and core genome MLST (cgMLST). To infer a neighbour-joining tree for phylogenetic analysis, Pathogenwatch calculates pairwise single nucleotide polymorphism (SNP) distances between genomes based on a concatenated alignment of 1972 core genes (Argimón et al., 2021). We then compared our genomes to those in Pathogenwatch, using cgMLST single linkage clustering, and selected those that had <10 allele differences (threshold=10) and <21 single-nucleotide polymorphisms (SNPs) (David et al., 2019) as the most closely related. All genome metadata in the neighbour-joining trees were added using iToL (Letunic and Bork, 2021).

Prediction of antibiotic resistance determinants (acquired and chromosomal mutations), and virulence traits (yersiniabactin-YbST, colibactin-CbST, aerobactin-AbST, salmochelin-SmST, and regulators of mucoid phenotype-RmpA, and RmpA2) was performed with Kleborate v2.3.0 (Lam et al., 2021). ABRicate v1.0.1 (https://github.com/tseemann/abricate) with *in-house* databases was used to detect additional *K. pneumoniae* virulence genes from BIGSdb-Pasteur (https://bigsdb.pasteur.fr/klebsiella/), and the chaperone-usher pili system (*kpiABCDEFG*) (Gato et al., 2020). The BacMet2 database, which is experimentally confirmed and available from http://bacmet.biomedicine.gu.se/, was manually curated to exclusively include proteins found in *Klebsiella pneumoniae* (including AcrAB-TolC, CpxAR, EmmdR, EmrDE, KdeA, KexD, KmrA, KpnEFO, MdtKM, OqxAB, PhoPQ, QacEΔ1, QacF, SugE, and TehA). This database, along with an *in-house* database of acquired metal tolerance proteins (including ArsR1H-ArsD1A1A2-ArsCBA3D2R2, PcoGE1ABCDRSE2, SilESRCFBAGP, MerRTPCADE, and TerZABCDEF-TerWY1XY2Y3) were used as reference for identifying biocide (QACs) and metal tolerance, respectively. The BLASTX 2.14.0+ (Camacho et al., 2009), with additional parameters “-evalue 0.001 -query_gencode 11”, was employed to perform the sequence alignment of the bacterial genomes against both databases. Confirmation of antibiotic-resistant novel alleles was performed using AMRFinderPlus v3.11.4 (Feldgarden et al., 2021).

Plasmid replicon typing was performed on all WGS-selected isolates using ABRicate v1.0.1 (https://github.com/tseemann/abricate) with PlasmidFinder database (from 2023-Jul-18) and pMLST v2.0 (Camacho et al., 2009; Carattoli et al., 2014) from the Centre for Genomic and Epidemiology (http://www.genomicepidemiology.org). IncFII_K_ plasmids were further characterized, as described by (Villa et al., 2010) (https://pubmlst.org/organisms/plasmid-mlst). We used the MOB-recon tool v3.1.0 from the MOB-suite package to confirm the location of the metal tolerance genes and reconstruct putative plasmids based on draft assemblies (Robertson and Nash, 2018; Robertson et al., 2020). Metal tolerance genes were considered part of a specific plasmid when identified by MOB-recon or when located on the same contig as the replicon/incompatibility determinant.

### 2.8 Statistical analysis

Differences in occurrence, antimicrobial resistance, and copper tolerance among *K. pneumoniae* P0, P1, P2, and P3 samples as well as isolates were analysed by Fisher’s exact test (α=0.05) using Prism software, version 8.1.1 (GraphPad).

## 3 Results

### 3.1 Occurrence of *K. pneumoniae* by poultry samples

We detected *K. pneumoniae* in 43% (n=24/56) of the samples collected throughout all steps of the poultry production chain. The highest occurrence was found in pre-slaughter faecal chicken samples (79%-11/14 flocks; in all but one farm), followed by derived chicken meat (50%-7/14 batches; all farms), and less frequently in one-day-old chicks (7%-1/14 flocks; one farm). Notably, *K. pneumoniae* was also detected in cleaned poultry houses (36%-5/14 flocks; four farms) (**Figure 2**). Although differences were observed between farms, a significant increase in occurrence was found between one-day-old and pre-slaughter chickens (p<0,05), and most (n=6/7) of these maintained *K. pneumoniae* carriage in chicken meat at the slaughter stage. Only three out of 11 pre-slaughter flocks carrying *K. pneumoniae* were raised in cleaned poultry houses that had previously tested positive for these bacteria (**Figure 2**).

**Figure 2.**
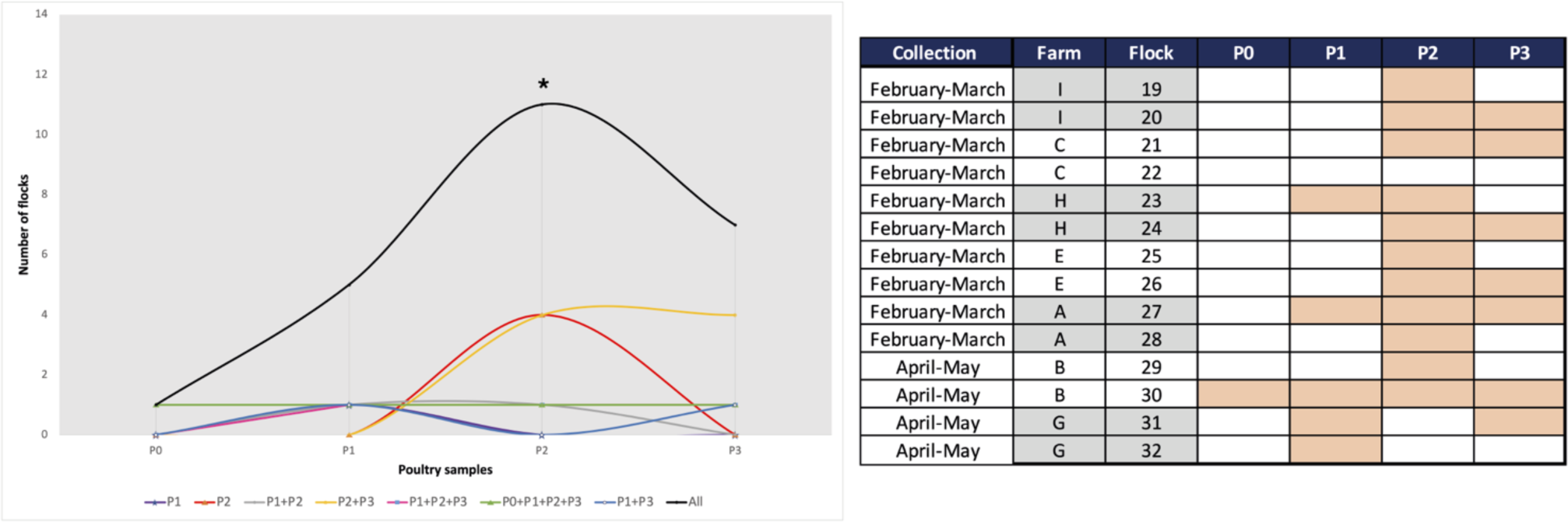
Occurrence of *K. pneumoniae* in poultry samples from the farm (P0, P1, and P2) and chicken meat (P3). *, P<0,05 (Fisher’s exact test) when comparing P0 with P2.

### 3.2 Diversity of *K. pneumoniae* along the poultry chain

We recovered 99 *K. pneumoniae* isolates from the 24 positive samples. Most isolates (>80%) were recovered from pre-slaughter chickens (65%-n=64/99, 11 flocks, all but one farm) and chicken meat (16%-n=16/99, 8 flocks, all farms) (**Figure 3**). With FT-IR, we were able to discriminate 24 putative KL-types, some of them identified in >4 isolates in the same (KL23, KL109) or in different samples (KL19, KL30, KL38, KL111). Furthermore, 54/99 (55%) isolates were correctly identified as carrying 14 KL-types included in the classification models used (KL3, KL10, KL14, KL19, KL21, KL23, KL28, KL30, KL38, KL64, KL102, KL106, KL109, KL111). Additionally, 15/99 (15%) isolates were grouped in 6 (n=2-4 isolates each) highly related spectral profiles. We used this information to select representative isolates by sample, antibiotic susceptibility profile and CuT *silA* gene for further characterization by *wzi* sequencing and/or whole genome sequencing.

**Figure 3.**
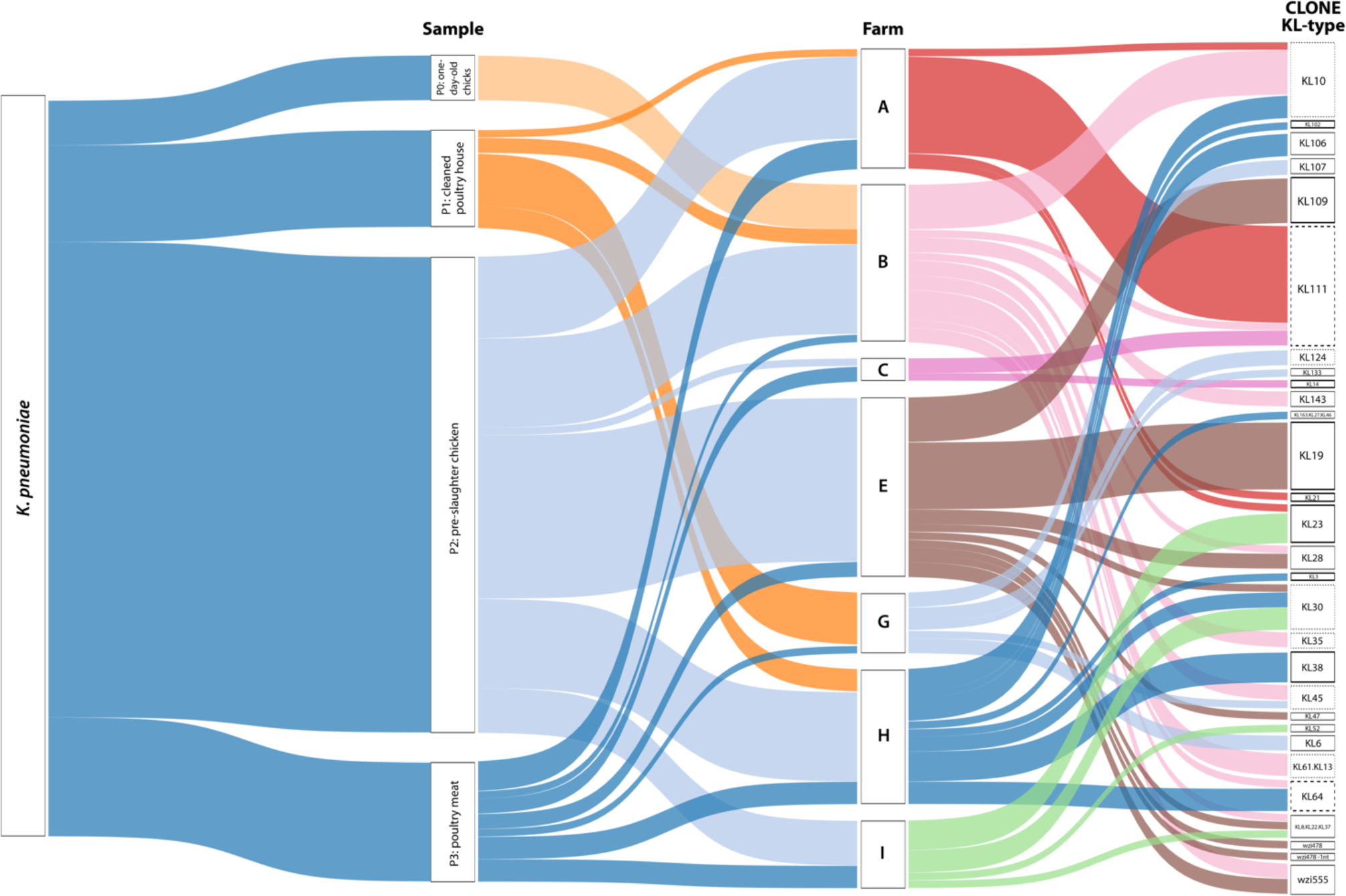
Sankey diagram representing, from left to right, the occurrence and diversity of *K. pneumoniae* by sample, farm, and KL-type. The width of each connection is proportional to the number of positive hits. The bold black line represents KL-types where 100% of the isolates were correctly identified by FTIR (KL3, KL14, KL19, KL21, KL23, KL38, KL102, KL109). The bold and dashed black line represents KL-types where at least 75% of the isolates were correctly identified by FTIR (KL64, KL111). The black dashed line represents KL-types where all the isolates exhibit highly related profiles recognized by FT-IR (KL10, KL30, KL35, KL45, KL124, KL61.KL13). The sensitivity and specificity of FTIR for KL-typing were 78% and 80%, respectively. The Sankey diagram was generated using Tableau Desktop 2023.3 (https://www.tableau.com/).

Considering FT-IR and sequencing data, a total of twenty-eight capsular locus (KL)-types were identified (13 detected in more than one sample), with the highest diversity observed in the pre-slaughter samples (**Figure 3**). The most frequent KL-types included KL111 (16%; P2+P3; 3 farms/4 flocks), KL10 (11%; P0+P2; 3 farms/3 flocks), KL19 (9%; P2; 1 farm/2 flocks), KL30 (6%; P2+P3; 3 farms/3 flocks), KL109 (6%; P2; 1 farm/1 flock) and KL23 (5%; P1+P2; 2 farms/2 flocks). Although less frequent, other KL types were shared by different farms, flocks, or stages (**Figure 3** and **Supplementary Table S1**).

### 3.3 Antibiotic susceptibility of *K. pneumoniae* recovered from poultry chain

More than 50% of the positive samples showed at least one *K. pneumoniae* isolate with decreased susceptibility to ciprofloxacin (83%-n=20/24) or resistance to nalidixic acid (58%-n=14/24), tetracycline (71%-n=17/24), sulphonamides (67%-n=16/24) or trimethoprim (63%-n=15/24) (**Figure 4).** Overall, 56% (n=55/99) of the isolates were MDR (**Supplementary Table S1**) identified in most samples (67%-n=16/24) from all farms. These were significantly more frequent in pre-slaughter (P2) than in chicken meat (P3) samples (p<0,05) (**Figure 4**). Approximately 20% of samples (including those from cleaned poultry houses, pre-slaughter chickens, and chicken meat), from diverse farms/flocks had at least one isolate resistant to extended-spectrum-cephalosporins (n=6 isolates/3 samples with an ESBL phenotype). Colistin-resistant isolates were identified only in one pre-slaughter flock sample (MIC>16 mg/L; *mcr* negative), all from the same clone KL109 recovered from SCAi+colistin plates. Most of the other isolates (n=91) were recovered using SCAi plate with (n=56) or without (n=35) an enrichment step.

**Figure 4.**
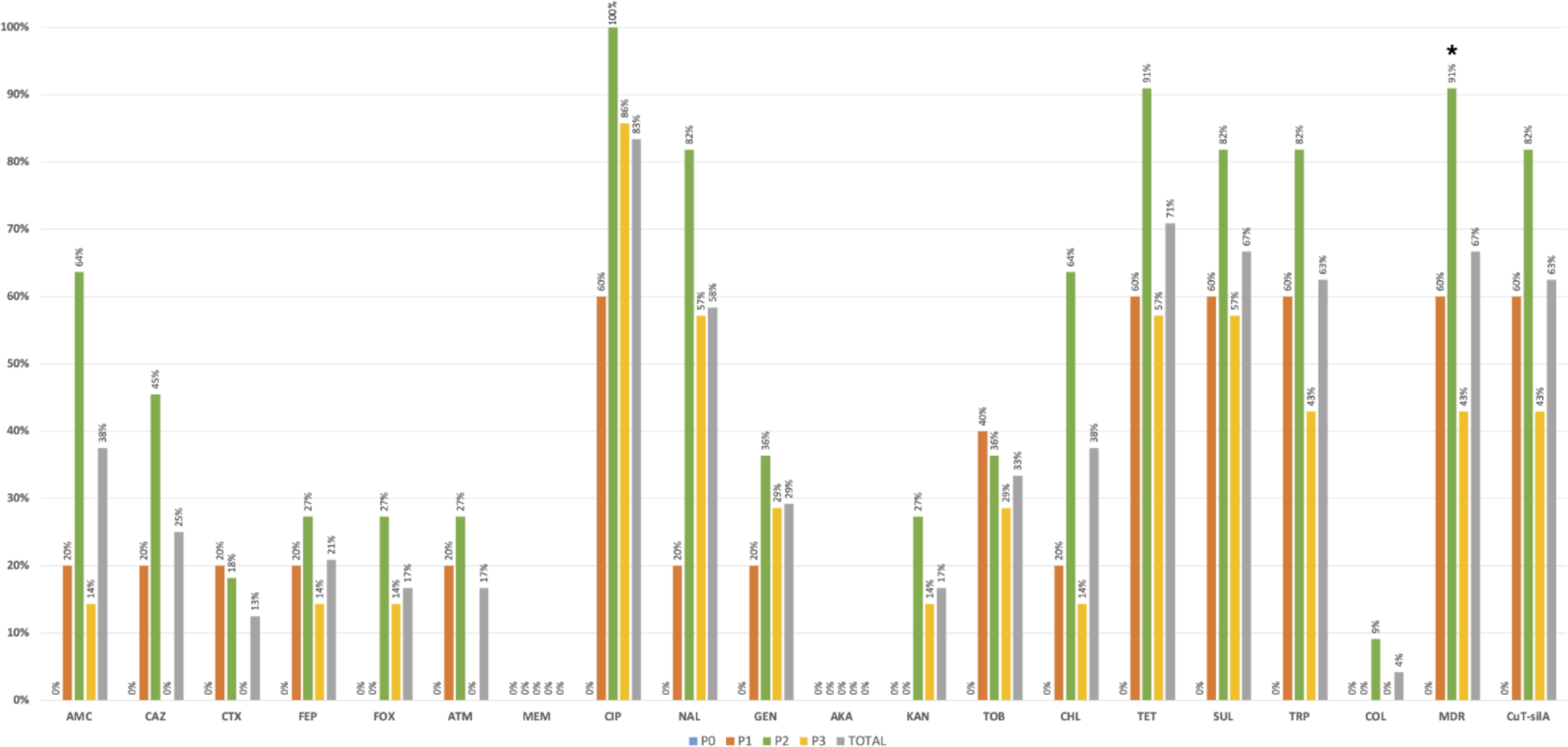
Occurrence of antibiotic-resistant *K. pneumoniae* in positive poultry samples at farm stages (P0, P1, and P2) and chicken meat (P3). *p<0,05 (Fisher exact test). Abbreviations: AMC, amoxicillin+clavulanic acid; CAZ, ceftazidime; CTX, cefotaxime; FEP, cefepime; FOX, cefoxitin; ATM, aztreonam; MEM, meropenem; CIP, ciprofloxacin; NAL, nalidixic acid; GEN, gentamicin; AKA, amikacin; KAN, kanamycin; TOB, tobramycin; CHL, chloramphenicol; TET, tetracycline; SUL, sulphonamides; TRP trimethoprim; COL, colistin; MDR, multidrug resistance; CuT-*silA*+, copper-tolerance *silA* gene.

### 3.4 Copper tolerance of *K. pneumoniae* recovered from poultry chain

The *silA* gene was observed in 52% (n=51/99) of *K. pneumoniae* isolates from all farms and most samples (63%, n=15/24), with similar rates between them (p>0,05) (**Figure 4).** More than 80% of *silA*-positive isolates were MDR (p<0.05) and were also found to be more resistant to amoxicillin+clavulanic acid, ciprofloxacin, nalidixic acid, tetracycline, sulphonamides, gentamicin, and trimethoprim than the *silA* negative ones (p<0.05) (**Figure 5**). Copper susceptibility assays were performed in 53% (n=52/99) of *K. pneumoniae* carrying or not copper tolerance genes representative of different farms, flocks, KL-types, and antibiotic resistance profiles (**Supplementary Table S1**). Phenotypic results were congruent with the genotype since all isolates with MIC_CuSO4_≥16mM carried the *silA* gene whereas those with MIC_CuSO4_<16mM did not (**Supplementary Table S1**).

**Figure 5.**
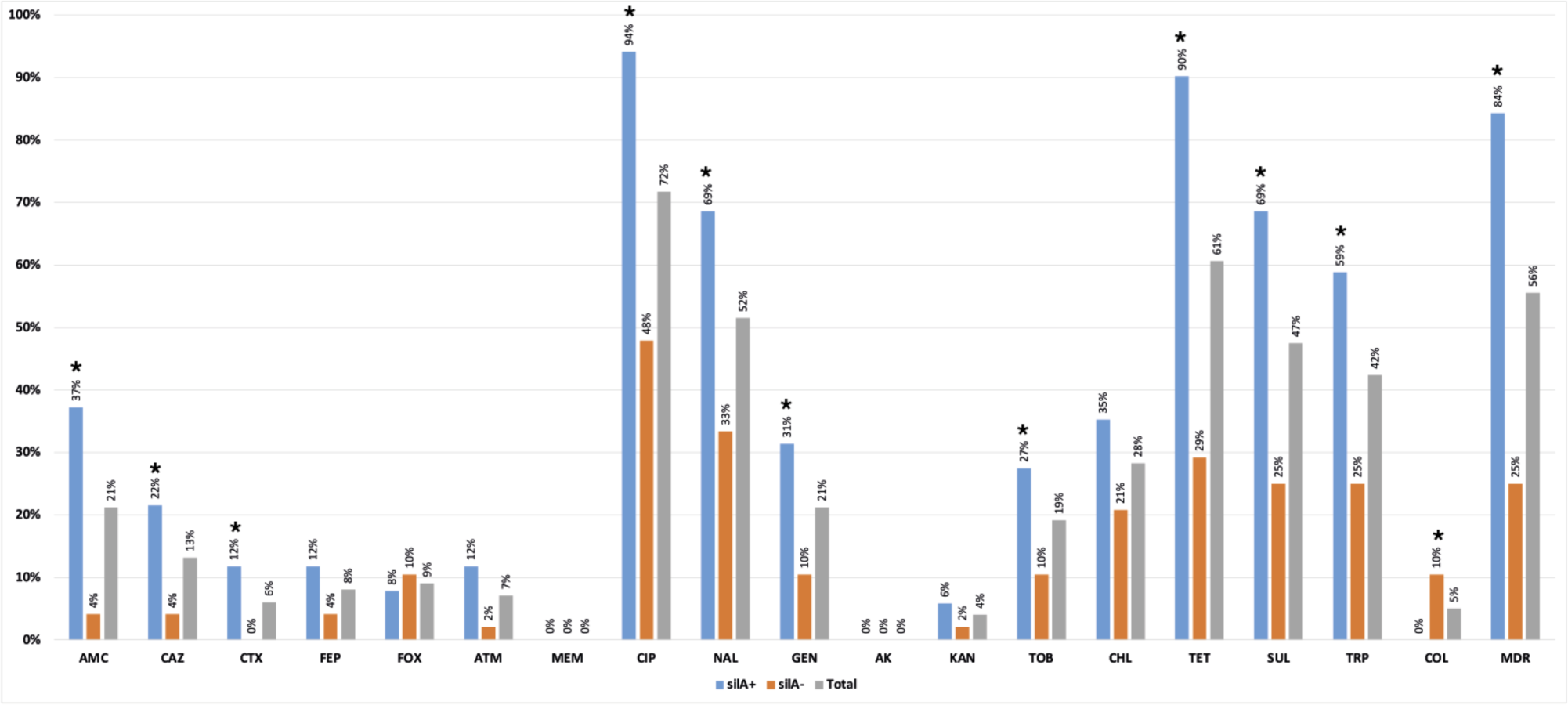
Percentage of antibiotic resistance detected among *silA*+ (n=51) and *silA*-(n=48) *K. pneumoniae* isolates. *p<0,05 (Fisher exact test). Abbreviations: AMC, amoxicillin+clavulanic acid; CAZ, ceftazidime; CTX, cefotaxime; FEP, cefepime; FOX, cefoxitin; ATM, aztreonam; MEM, meropenem; CIP, ciprofloxacin; NAL, nalidixic acid; GEN, gentamicin; AKA, amikacin; KAN, kanamycin; TOB, tobramycin; CHL, chloramphenicol; TET, tetracycline; SUL, sulphonamides; TRP trimethoprim; COL, colistin; MDR, multidrug resistance.

### 3.5 Benzalkonium chloride tolerance of *K. pneumoniae* recovered from poultry chain

The susceptibility to BZC was determined for 45 *K. pneumoniae* (all carrying genes previously associated with QAC tolerance), with diverse epidemiological and clonal backgrounds (**Supplementary Table S2**). The MIC_BZC_ ranged between 4-64 mg/L, with MIC_50_=16 mg/L and MIC_90_=32 mg/L. The highest MIC_BZC_ of 64 mg/L [non-wild-type using ECOFF of 32 mg/L proposed by (Morrissey et al., 2014)] was observed in the ST280-KL23 lineage from a pre-slaughter chicken sample. Ten QACs’ genotypes (19 to 24 genes each) were identified. A variable occurrence of *emrE* (n=1, 1 ST), *qacF* (n=7, 5 STs), *qacEΔ1* (n=22, 14 STs); *kexD* (n=37, 25 STs); *oqxAB* (n=43, 30 STs) and *kmrA* (n=44, 30 STs) was found, with the remaining genes detected in 100% of the genomes (**Figure 6**). However, no differences were observed between MIC distribution and clones, sources, MDR phenotype or the presence or absence of genes previously associated with QAC tolerance, including *qac* genes (**Figure 6 and Supplementary Table S2**). The only exception was the ST6552-KL109 isolate lacking *kmrA*, coding for an efflux pump of the Major Facilitator Superfamily, and presenting MIC_BZC_ of 4 mg/L. The MBC_BZC_ for all tested isolates was the same as the MIC_BZC_. *K. pneumoniae* showing the highest MBC_BZC_ of 32-64 mg/L comprised isolates from diverse farms, sources, and clones, including one of the most prevalent in the present study (ST11-KL106, farm H).

**Figure 6.**
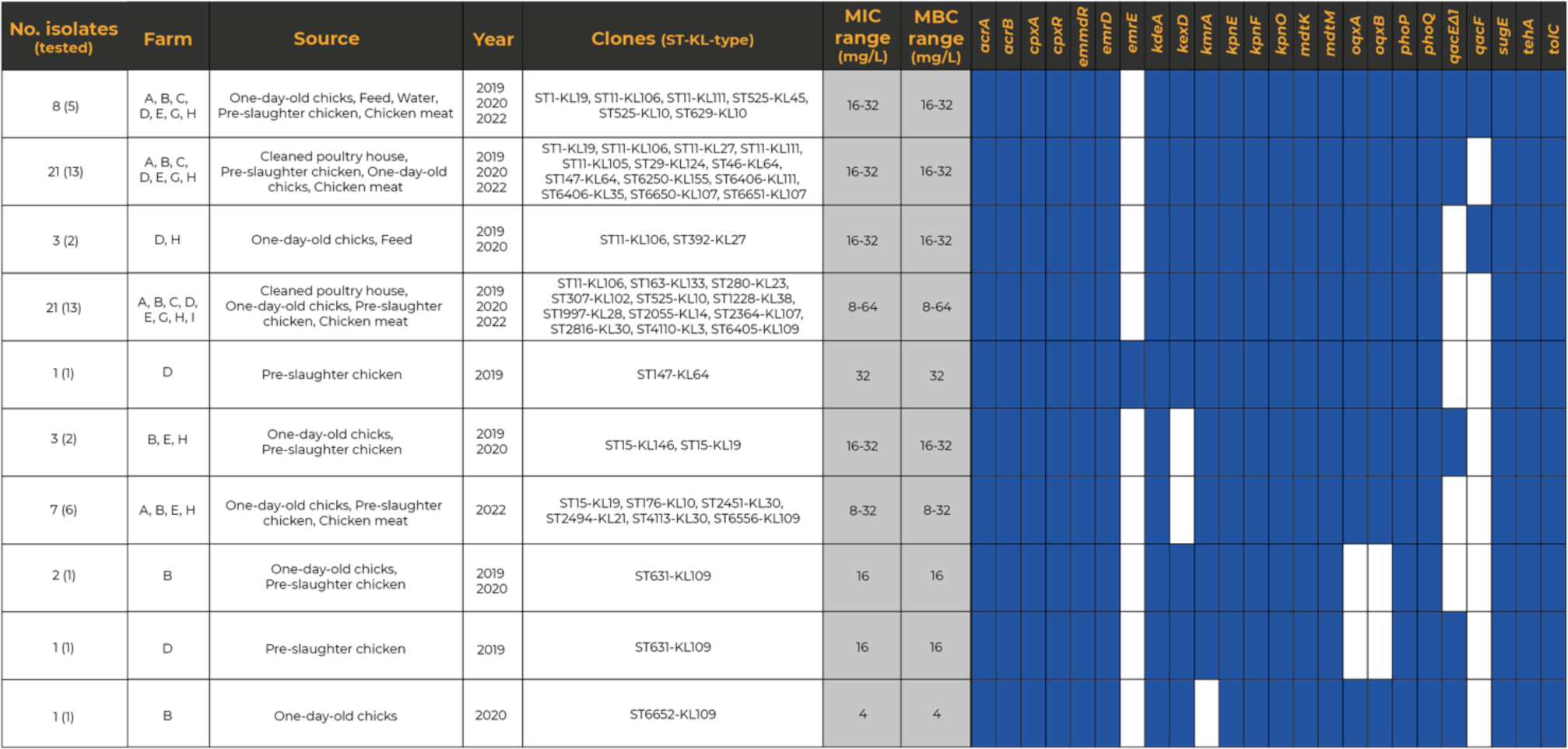
Heatmap representing the distribution of QAC tolerance genes across representative *Klebsiella pneumoniae* isolates (n=45). The presence of a specific QAC tolerance gene is indicated by a filled blue square.

### 3.6 Whole-genome analysis of *K. pneumoniae* poultry-associated isolates

The genetic relatedness of *K. pneumoniae* isolates revealed a clonal diverse population from 2019 to 2022 [this collection and (Mourão et al., 2023)]. Overall, the isolates were assigned to 31 STs (7 new STs) and 37 lineages, including the globally successful ST11-KL106, ST11-KL111, ST15-KL19, ST147-KL64, ST280-KL23, and ST307-KL102 (**Figure 7** and **Supplementary Table S2**). Based on the cgMLST analysis and the proposed thresholds (<10 allele differences), 15 clusters (differing by 0-207 SNPs) were identified among poultry *K. pneumoniae* (**Figure 7** and **Supplementary Table S3**). Those clusters included isolates from different sample types from the same farm: pre-slaughter chicken faeces (ST15-KL19), water and chicken (ST11-KL106) pre-slaughter chicken faeces and cleaned poultry houses (ST1228-KL38; ST6651-KL107), pre-slaughter chicken faeces and derived meat (ST1-KL19; ST11-KL111; ST6405-KL109), or cleaned poultry-house and chicken meat (ST29-KL124). Furthermore, some lineages (e.g., ST631-KL109, ST6651-KL107, ST6406-CG147-KL111) were detected over time (**Figure 7**).

**Figure 7.**
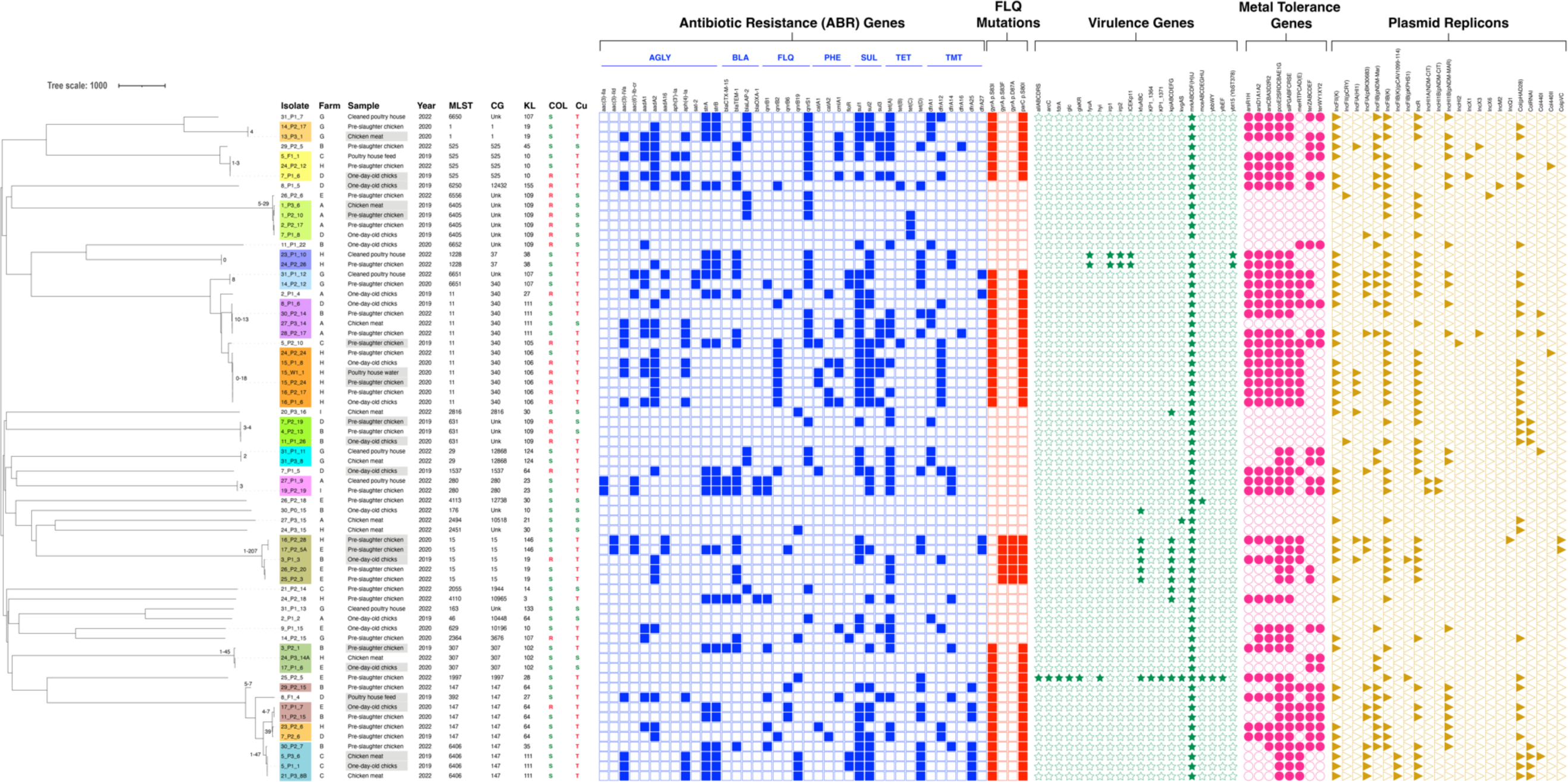
A neighbour-joining tree representing the phylogenetic relationships among the 68 *K. pneumoniae* genomes recovered between 2019-2022 was constructed from the Pathogenwatch pairwise-distance matrix [i.e., based on single nucleotide polymorphisms (SNPs) called in 1972 core genes]. Scale bar units represent substitutions per variant site. The SNPs among our isolates from the 15 main clusters (< 10 alleles difference) are represented in each branch. The isolates exhibiting grey shading correspond to the ones already published by (Mourão et al., 2023). The associated metadata for all isolates was added using iTOL (https://itol.embl.de/). Each coloured-filled shape represents the presence of relevant antibiotic resistance, virulence-associated, metal-tolerance genes, and plasmid replicons associated with well-defined incompatibility groups. Only known mutations conferring fluoroquinolone resistance are presented. *Klebsiella* intrinsic antibiotic resistance (*bla*_SHV-1_, *bla*_SHV-11_, *bla*_SHV-26_, *bla*_SHV-27_, *bla*_SHV-28_, *bla*_SHV-38_, *bla*_SHV-142_, *bla*_SHV-187_, *fosA*, *oxqAB*) and metal tolerance (*arsBCR*, *cusABFCRS*) genes were not represented. Abbreviations: AGLY, aminoglycosides; BLA, β-lactams; CG, Clonal Group; FLQ, fluoroquinolones; KL, K-Locus; MLST, Multilocus Sequence Typing; PHE, phenicols; SUL, sulphonamides; TET, tetracycline; TMT, trimethoprim; Unk, unknown.

To investigate the phylogenetic relationship of our genomes with others, we conducted a thorough analysis using Pathogenwatch, incorporating diverse sources, regions, and timeframes (**Figure 8; Supplementary Figure S1**). The analysis revealed less than 10 alleles differences in 18 lineages, with ST11-KL27, ST11-KL111, ST15-KL19, ST15-KL146, ST280-KL23, ST392-KL27, ST1537-KL64 and ST1997-KL28 with fewer than 21 single nucleotide polymorphisms (SNPs) (a threshold recently proposed for *K. pneumoniae* transmission in healthcare settings) (David et al., 2019) when compared to genomes from various sources (**Figure 8**). Most were linked to genomes from human infections in diverse EU countries, including Portugal (ST1997-KL28) (**Figure 8**).

**Figure 8.**
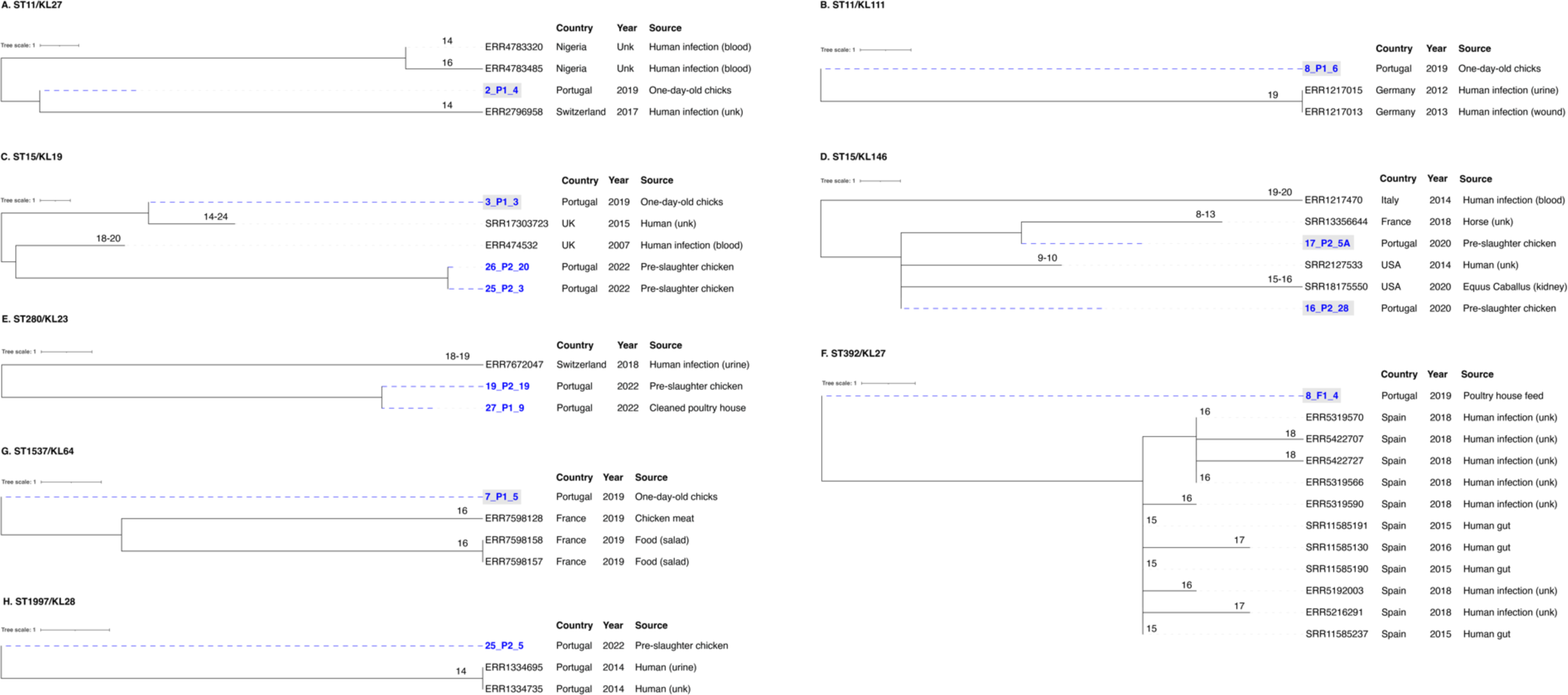
Neighbour-joining trees representing the phylogenetic relationships among the 68 *K. pneumoniae* genomes recovered between 2019-2022 and those available in Pathogenwatch with less than 21 SNPs. **(A)** ST11-KL27, **(B)** ST11-KL111, **(C)** ST15-KL19, **(D)** ST15-KL146, **(E)** ST392-KL27, **(F)** ST1537-KL64, **(G)** ST280-KL23, and **(H)** ST1997-KL28. The genome selection was performed using the cgMLST single linkage clustering to include the ones with less than 10 allele differences (threshold=10)*. Then these genomes were used to infer a neighbour-joining tree from the Pathogenwatch pairwise-distance matrix (i.e., based on single nucleotide polymorphisms-SNPs called in 1972 core genes). Scale bar units represent substitutions per variant site. The number of substitutions between our isolates and the ones available in Pathogenwatch are represented in each branch. All isolates’ associated metadata (country, collection date, and source of isolation) was added using iTOL (https://itol.embl.de/). \**K. pneumoniae* isolates from clonal lineages with <10 allele differences include: 2019-2020 collection - ST1-KL19, ST11-KL105, ST11-KL106, ST11-KL27, ST11-KL111, ST15-KL19, ST15-KL146, ST147-KL64, ST307-KL102, ST392-KL27, ST1537-KL64, and ST6250-KL155 (1); 2022 collection - ST11-KL106, ST11-KL111, ST15-KL19, ST147-KL64, ST280-KL23, ST307-KL102, ST525-KL45, ST1228-KL38, ST2055-KL14, ST1997-KL28, ST4110-KL3. The isolates exhibiting grey shading correspond to the ones already published by (Mourão et al., 2023).

The WGS revealed a high load and diversity in antibiotic resistance genes between 2019-2022 (**Figure 7** and **Supplementary Table S2**). Thirty-eight acquired genes (7 classes) were detected, including clinically-relevant ones (*bla*_CTX-M-15_, *qnrB* and *qnrS*). More than 75% (n=51/68) of the genomes carried genes coding for ≥ 3 classes of antibiotics, with the most frequent including aminoglycosides (*strA*/*strB/aadA*), sulphonamides (*sul1/sul2*), tetracyclines (*tet(A)*/*tet(D)*), trimethoprim (*dfrA*) and phenicols (*catA, cmlA*) (**Figure 7** and **Supplementary Table S2**). The most frequent determinant of resistance to ciprofloxacin was the presence of chromosomal mutations in the topoisomerase genes *gyrA* and *parC* (59%-n=40/68) over 2019-2022. The ParC S80I mutation was present in all genomes, independently of the clonal lineage. The single GyrA S83I mutation was observed in all genomes except for ST15, which had two others simultaneous *gyrA* mutations (S83F and D87A). Furthermore, we noticed a decrease in isolates carrying chromosomal mutations implicated in colistin resistance between 2019-2020 (12 lineages) and 2022 (1 lineage ST6556/KL109) (**Figure 7** and **Supplementary Table S1**). Concerning the virulence genes, all but one isolate carried the chromosomally encoded fimbriae cluster (*mrkABCDFHIJ*). The worldwide-disseminated high-risk clone ST15 isolates carried the ferric uptake (*kfuABC*) and the chaperone-usher pili systems (*kpiABCDEFG*) whereas ST1228 isolates the yersiniabactin siderophore (*ybt*15-YbST378/ICEKp11) in addition to the genes encoding for proteins required for yersiniabactin (*irp1*, *irp2*) and siderophores (*fyuA* gene) (**Figure 7**).

### 3.7 Whole-genome analysis of metal tolerance genés occurrence and location

Metal tolerance genes were observed in most (75%, n=51/68) isolates, specifically copper/silver (*sil*/*pco*), arsenic (*ars*), mercury (*mer*), and/or tellurite (*ter*) (**Figure 7**). Notably, the most frequent *sil* and *pco* cluster (69%, n=47/68) was frequently found alongside the *ars* (77%, n=36/47), *ter* (51%, n=24/47) and/or *mer* (49%, n=23/47) operons. We detected, on average, four plasmids per genome (ranging from 1 to 10) and 25 distinct plasmid incompatibility groups (with 1-8 replicon types per genome) (**Supplementary Table S2**). The most prevalent groups were IncFIB_K_ (65%, n=44/68), followed by IncFII_K_ (59%, n=40/68), and IncR (56%, n=38/68), often in different combinations with FIA, other FIB types (such as FIB_pNDM-MAR_, FIB_pCAV1099_, FIB_pKPHS1_), and/or HI1B (**Figure 7** and **Supplementary Table S2**). Col plasmids were also frequently observed (54%, n=37/68 genomes), whereas other classical incompatibility groups (IncHI2, IncX, IncQ, IncM) were infrequent.

Copper tolerance genes *sil* and *pco* were plasmid located in all but one isolate (98%-n=46/47) primarily within IncFIB_K_+IncFII_K_ plasmids (51%, n=24/47, approximately 80-350 Kb) of different IncFII_K_ groups based on pMLST (2, 4, 5, 7, 8, 21-like). Additionally, copper tolerance genes were co-localized with variable genes coding for resistance to aminoglycosides (*aac(3)-IIa*, *aac(3)-Iva*, *aac(6’)-Ib-cr*, *aph(4)-Ia*, *aadA1*, *aadA2*, *aadA16*, *strA*-*strB*), fluoroquinolones (*qnrB1*, *qnrB2*, *qnrB6*, *qnrS1*), chloramphenicol (*catA1*, *catA2*, *cmlA1*), sulphonamides (*sul1, sul2*, *sul3*), tetracycline (*tetA*, *tetD*), trimethoprim (*dfrA12*, *dfrA14*, *dfrA25*, *dfrA27*) and β-lactams (*bla*_TEM_, *bla*_CTX-M-15_, *bla*_OXA-1_) (**Figure 7** and **Supplementary Table S4**). Co-location with *qac* tolerance genes (*qacEΔ1* and/or *qacE*, *qacF*) was also observed in six isolates. These plasmids carrying *sil* and *pco* were like others described in humans across multiple countries (**Supplementary Table S4**), occasionally harbouring *bla*_CTX-M-15_, *bla*_DHA-1_, *bla*_NDM-1,_ *bla*_OXA-1_, *bla*_SHV-30_, and/or *qnr* genes.

## 4 Discussion

This is a distinctive study that explores *K. pneumoniae* occurrence and diversity throughout the entire chicken farm-to-fork chain from a long-term perspective. It is also one of the few studies that offer evidence that poultry serves as a reservoir and source of *K. pneumoniae* strains with clinically relevant features, including genes coding for antibiotic resistance, metals, and/or biocide tolerance, while the identification of clones identical to those in clinical settings further supports *K. pneumoniae* as a foodborne pathogen.

Our data strongly emphasize that intensively farmed chicken production and their meat are relevant sources of *K. pneumoniae* and antibiotic-resistant isolates. This remains true even after EU veterinary antimicrobial sales decreased by 46.5% between 2011 and 2021 (European Medicines Agency, 2022). During the studied period [this study and (Mourão et al., 2023)], resistance remained high to antibiotics commonly used in poultry, such as tetracycline, sulphonamides, and/or quinolones (Alliance to Save our Antibiotics, 2016; Gržinić et al., 2023). However, resistance to colistin, a critically important antibiotic for treating human infections caused by Gram-negative MDR strains, has significantly decreased from 61% in 2019-2020 samples to 4% in 2022. This reduction suggests that the efforts of the long-term ban on colistin use in food-animal production (>4 years) have yielded promising results not only by limiting *mcr* (Ribeiro et al., 2021) but also other colistin-resistance genotypes (Mourão et al., 2023).

In conventional chicken production farms, diverse antimicrobial treatments are commonly administered throughout the fattening period, with differences by country and region (Joosten et al., 2019; Kasabova et al., 2021). In this study, all but one flock received antibiotics during the 26-43 days production cycle, which may explain the higher occurrence of MDR *K. pneumoniae* in pre-slaughter chickens compared to samples from previous stages. Furthermore, the contribution of poultry house management practices like inadequate disinfection or short vacancy periods between flocks, cannot be excluded (Daehre et al., 2018; Zhai et al., 2020; Franklin-Alming et al., 2021; Kaspersen et al., 2023). Even with standard cleaning and disinfection protocols across all the studied farms, a wide variety of *K. pneumoniae* strains, including MDR and clinically relevant lineages, can persist within the farm environment and/or until the meat becomes available to consumers. We found closely related MDR *K. pneumoniae* isolates in both cleaned poultry houses and pre-slaughter chickens (ST1228-KL38, ST29-KL24, ST280-KL23) from the same or different flocks or farms. The idea that strains re-introduction might come from parent flocks seems less likely since we observed minimal contamination (1 flock) during the reception of one-day-old chickens. This study also highlights water and feed as sources of MDR *K. pneumoniae* lineages (ST525-KL10, ST11-KL106), but at lower rates than chicken faeces, as observed in (Mourão et al., 2023). Noteworthy, some *K. pneumoniae* lineages (such as ST1-KL19, ST11-KL111, ST6405-KL109, and ST6406-CG147-KL111) persist throughout the chickens’ lifecycle until their meat becomes available to consumers. Additionally, cross-contamination at slaughterhouses is also plausible given the identification of *K. pneumoniae* lineages in chicken meat samples that were not traced back to their originating farm. The higher contamination compared with the previous study (43-50% vs 17%) and the diversity of *K. pneumoniae* strains (Mourão et al., 2023), might be attributed to the cultural methodology employed (antibiotic-free selection using SCAi medium with and without BPW enrichment). This approach was specifically designed to enhance the investigation of the inherent *K. pneumoniae* populations, as previously reported (Rodrigues et al., 2022). FT-IR proved to be a reliable approach to assessing the clonal relationship between *K. pneumoniae* isolates from poultry origin, being able to correctly identify closely related isolates (representing 70% of the sample) and KL-types (78% predicted by the machine-learning models) and discard unrelated isolates or KL-types not represented by the models used, as happened for isolates from human origin (Novais et al., 2023a). The use of this technique represented a useful tool to quickly identify common isolates from the same or different samples and choose representative isolates to sequence by WGS, reducing the cost and time associated with typing.

In our comparative genomic analysis using WGS, we identified closely related MDR *K. pneumoniae* lineages such as ST15-KL19, ST11-KL111, ST280-KL23, and ST1997-KL28 shared between poultry and human clinical isolates, even when applying strict criteria of less than 21 SNPs (David et al., 2019). Additionally, certain poultry strains shared a substantial repertoire of accessory genes, including fluoroquinolone resistance mutations and/or virulence genes with clinically relevant human clones (e.g., ST15, ST147, ST307) identified in previous studies (Peirano et al., 2020; Rodrigues et al., 2022, 2023). This data strengthens the argument for the possible transmission of these strains from food animals to humans (Büdel et al., 2020; Rodrigues et al., 2022; Thorpe et al., 2022; Crippa et al., 2023; Kaspersen et al., 2023; Zou et al., 2023), solidifying poultry’s position as both a reservoir and source of globally dispersed, clinically relevant *K. pneumoniae* lineages (Mourão et al., 2023).

The persistence of MDR *K. pneumoniae* lineages within the poultry chain [this study, (Kaspersen et al., 2023; Mourão et al., 2023)], suggests the presence of adaptive environmental factors beyond antibiotic resistance. We observed elevated rates of *K. pneumoniae* carrying *sil* and *pco* operons, along with a copper tolerance phenotype (>16 mM) in poultry samples. This suggests that the incorporation of copper-supplemented chicken feed might contribute to the selection of copper-tolerant and MDR strains within such production environments. However, earlier research, whether grounded in wet lab experiments or mathematical models, has indicated that the concentrations of copper necessary to foster the emergence of copper-tolerant bacteria might be significantly below their corresponding MIC values (Gullberg et al., 2014; Arya et al., 2021). Furthermore, the coexistence of various pollutants appears to further lower the minimum selective concentration estimates for individual antimicrobials (Gullberg et al., 2014; Arya et al., 2021). The *sil* and *pco* operons are often located on plasmids that carry an array of other metal, *qac*, metabolic, and/or specific antibiotic resistance genes, fostering their persistence within the environment through co-selection events driven by antimicrobial usage, coupled with the frequent presence of mechanisms that prevent plasmid loss.

This study also reveals a high abundance of QACs tolerance genes in *K. pneumoniae* lineages, although they do not appear to correlate with phenotype, as observed in other studies regardless of testing methods (Abuzaid et al., 2012; Morrissey et al., 2014; Wu et al., 2015; Vijayakumar et al., 2018; Gual-de-Torrella et al., 2022). Nevertheless, our WGS approach identified a wide range of QAC tolerance genes, underscoring the pressing need for establishing reliable genotypic-phenotypic correlations to elucidate QAC tolerance mechanisms (Hipólito et al., 2023). While several studies have reported that bacterial strains with an elevated MIC or MBC remained susceptible to the in-use BZC concentration (Vijayakumar et al., 2018; Maillard, 2022), it is essential to comprehend the environmental factors (e.g., temperature, pH) associated with the expression of these genes (Gual-de-Torrella et al., 2022; Pereira et al., 2023) to clarify the environmental, clinical, or veterinary/industrial implications of bacteria with a reduced biocide susceptibility. Further studies are needed to investigate the impact of inappropriate biocide usage or low concentrations, which can act as stressors without killing bacterial pathogens, potentially promoting antimicrobial resistance, and facilitating the transfer of antimicrobial resistance genes (Maillard, 2022; Maillard and Pascoe, 2023).

## 5 Conclusions

Our study reveals chicken production as a significant reservoir hosting a diverse range of clinically relevant *K. pneumoniae* clones, including MDR, copper-tolerant and enriched in QAC tolerance genes. The identification of clones identical to those in clinical settings supports *K. pneumoniae* as a foodborne pathogen. Various sources of contamination (such as feed, water, poultry houses, and cross-contamination) contribute to the persistence of *K. pneumoniae* throughout the production chain, emphasizing that, despite a decrease in its occurrence, certain clones still reach chicken meat even with implemented safety measures in place.

The co-occurrence of copper and/or QAC tolerance genes on highly prevalent MDR plasmids suggests that these have been circulating in various *K. pneumoniae* populations and phenotypic validation (at least in the case of copper) supports the possibility that these genes may play a role in the co-selection of these plasmids or strains under certain conditions within the food production chain or other environmental settings.

Further studies are needed to assess the implications of these *K. pneumoniae* lineages on food safety and the risk of transmitting antibiotic resistance to humans. Additional studies are imperative to elucidate the external factors (such as environmental conditions) that drive *K. pneumoniae*’s adaptation towards antimicrobial resistance. Addressing these complexities can contribute to the development of effective strategies to safeguard animal welfare, enhance food safety, and mitigate public health risks associated with clinically-relevant *K. pneumoniae* lineages and antibiotic resistance.

## 6 Conflict of Interest

*The authors declare that the research was conducted in the absence of any commercial or financial relationships that could be construed as a potential conflict of interest*.

## 7 Ethics statement

Ethical review and approval were not required because the samples were taken from farm environmental points (e.g., boxes, floor) as well as after routine slaughter, under the auspices of the poultry farm company, which includes oversight by the veterinary team.

## 8 Author Contributions

**Joana Mourão:** Writing – original draft, Writing – review & editing, Data Curation, Formal Analysis, Investigation, Methodology, Software, Visualization; **Mafalda Magalhães:** Writing – review & editing, Investigation, Methodology; **Marisa Ribeiro-Almeida:** Writing – review & editing, Formal Analysis, Investigation, Methodology; **Andreia Rebelo:** Writing – review & editing, Investigation, Visualization; **Carla Novais:** Writing – review & editing, Conceptualization, Formal Analysis, Funding acquisition, Investigation; **Luisa Peixe:** Writing – review & editing, Funding acquisition; **Ângela Novais:** Writing – review & editing, Conceptualization, Formal Analysis, Methodology; **Patrícia Antunes:** Writing – original draft, Writing – review & editing, Conceptualization, Formal Analysis, Funding acquisition, Investigation, Methodology, Project administration, Software, Supervision.

## 9 Funding

This work was supported by the Applied Molecular Biosciences Unit - UCIBIO which is financed by national funds from FCT - Fundação para a Ciência e a Tecnologia [UIDP/04378/2020 and UIDB/04378/2020], by the Associate Laboratory Institute for Health and Bioeconomy-i4HB [LA/P/0140/2020], by the AgriFood XXI I&D&I project [NORTE-01-0145-FEDER-000041] co-financed by the European Regional Development Fund (ERDF) through NORTE 2020 (Programa Operacional Regional do Norte 2014/2020) and by the University of Porto and Soja de Portugal [Grant N° IJUP-Empresas-2021-14]. Joana Mourão and Ângela Novais were supported by national funds through FCT/MCTES in the context of the Scientific Employment Stimulus (2021.03416.CEECIND and 2021.02252.CEECIND, respectively). Marisa Ribeiro-Almeida and Andreia Rebelo were supported by PhD fellowships from FCT (SFRH/BD/146405/2019 and SFRH/BD/137100/2018, respectively). The authors are greatly indebted to all the financing sources.

## 10 Acknowledgements

We thank the Institut Pasteur teams for the curation and maintenance of BIGSdb-Pasteur databases at http://bigsdb.pasteur.fr/. We also thank MSc Sofia Ribeiro for the Figure 1 design, the students António Magalhães (FCNAUP), Francisca Pereira (FCNAUP), and Cátia Matos (FFUP) for technical support, and the staff of the participating farms and slaughterhouses for their kind cooperation.

## 12 Supplementary Material

The Supplementary Material for this article can be found online.

## 13 Data Availability Statement

The sequencing data of the 48 isolates produced during this study are accessible in the European Nucleotide Archive (ENA) (https://www.ebi.ac.uk/ena). These data are stored under the Bioproject accession number PRJEB62836, with the specific accession numbers for each isolate detailed in **Supplementary Table 2**.

## Notes

### Competing Interest Statement

The authors have declared no competing interest.

