## Supplementary figures and images for "Decoding *Klebsiella pneumoniae* in Poultry Chain: Unveiling Genetic Landscape, Antibiotic Resistance, and Biocide Tolerance in Non-Clinical Reservoirs"

### Supplemental Figure S1

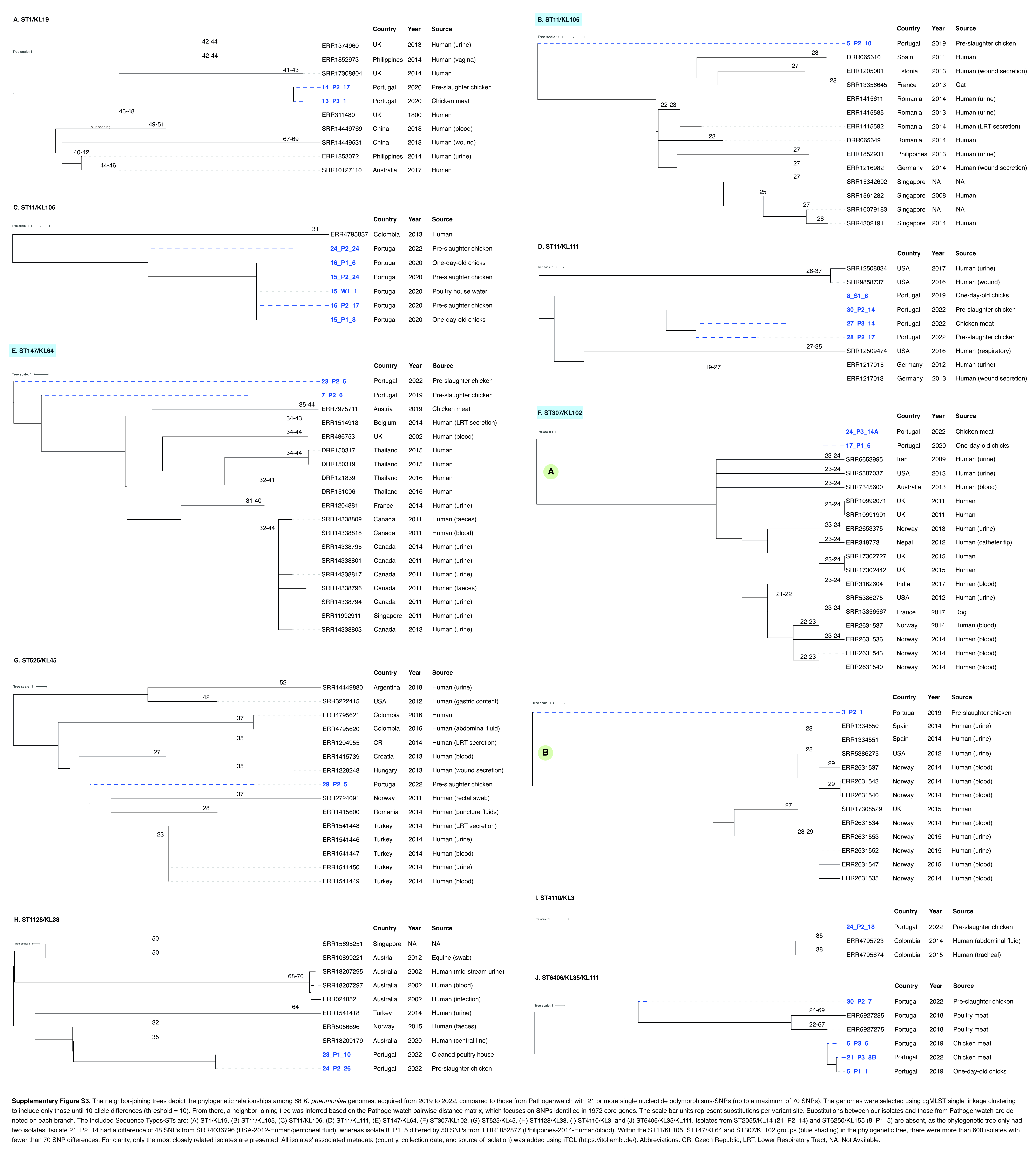
